# Protein Function Prediction from Three-Dimensional Feature Representations Using Space-Filling Curves

**DOI:** 10.1101/2022.06.14.496158

**Authors:** Dmitrij Rappoport, Adrian Jinich

**Affiliations:** Department of Chemistry, University of California, Irvine, 1102 Natural Sciences 2, Irvine CA, 92697; Weill Cornell Medicine, 1300 York Ave, Box 65, New York, NY 10065

## Abstract

Compact and interpretable structural feature representations are required for accurately predicting properties and the function of proteins. In this work, we construct and evaluate three-dimensional feature representations of protein structures based on space-filling curves. We focus on the problem of enzyme substrate prediction, using two ubiquitous enzyme families as case studies: the short-chain dehydrogenase/reductases (SDRs) and the S-adenosylmethionine dependent methyltransferases (SAM-MTases). Space-filling curves such as Hilbert curve and the Morton curve generate a reversible mapping from discretized three-dimensional to one-dimensional representations and thus help to encode three-dimensional molecular structures in a system-independent way and with a minimal number of parameters. Using three-dimensional structures of SDRs and SAM-MTases generated using AlphaFold2, we assess the performance of the SFC-based feature representations in predictions on a new benchmark database of enzyme classification tasks including their cofactor and substrate selectivity. Gradient-boosted tree classifiers yield binary prediction accuracy of 0.766–0.906 and AUC (area under curve) parameters of 0.828–0.922 for the classification tasks. We investigate the effects of amino acid encoding, spatial orientation, and (the few) parameters of SFC-based encodings on the accuracy of the predictions. Our results suggest that geometry-based approaches such as SFCs are promising for generating protein structural representations and are complementary to the highly parametric methods, for example, convolutional neural networks (CNNs).

## Introduction

The function of the vast majority of proteins in general, and enzymes in particular, remains unknown. For example, according to the UniProt Knowledgebase (UniProtKB),^1^ ≈ 60% of human proteins have the lowest annotation category (1-out-of-5), while only ≈ 3% have functional annotations with experimentally evidence that merit the highest annotation score.^2^ While recent advances in protein structure prediction ^3,4^ will accelerate protein functional annotation, how best to map structure to function remains a challenging and open problem.

Different representations of protein structure can be used to map structure to function. One common approach uses 3D convolutional neural networks.^5–7^ Concentric shells surrounding specific protein sites can also be used to represent the spatial distribution of biochemical properties.^8^ Other methods use graph representations of enzymes and their active sites.^9,10^ For example, Gligorijević et al.^9^ generate amino contact maps from protein structures, and use the resulting adjacency matrix as input for a graph convolutional network. The distribution of torsion angles and pairwise distances, extracted for each amino acid type separately, can also generate feature maps for downstream protein or enzyme functional prediction.^7,11^

A specific challenge in predicting function from protein structure is incorporating the full three-dimensional structure in addition to the amino acid sequence and the topological properties, as expressed, for example, by the residue contact maps.^12–17^ Two possible paths towards this goal are using minimally processed structural data together with highly tuned machine learning models such as 3D convolutional neural networks, or using optimized feature representations in combination with simpler predictive models. Here we pursue the second option with the expectation that the results should generalize well and should be easy to interpret. We thus seek feature representations of molecular three-dimensional structures that are reversible, lend themselves to machine learning algorithms, allow for comparisons between structures with different numbers of atoms, and incorporate the invariances of molecular structures such as rigid translations, rotations, and atom renumbering.

In this paper we explore the use of space-filling curves (SFCs)^18,19^ for generating discrete, invariant, and reversible feature representations of three-dimensional protein structures. SFCs were discovered in the late nineteenth century and describe a class of curves in *d*-dimensional Euclidean space ℝ^*d*^ (*d* ≥ 2) with the property that they pass through every point of a square (for *d* = 2) or cube (for *d* = 3). Specifically, a SFC is defined by a continuous mapping of the (one-dimensional) unit interval [0, 1] onto an *n*-dimensional object with non-zero area (for *d* = 2), volume (for *d* = 3), or its higher-dimensional equivalent (for *d* > 3). The SFC thus “linearizes” the set of points in ℝ^*d*^ (*d* ≥ 2) by prescribing a well-defined order of their traversal. The first SFC was described by Peano in 1890.^20^ In the following year, Hilbert published the definition and the geometric procedure for generating the eponymous curve.^21^ The Morton curve (also named Z-order curve) first appeared in a technical report in 1966.^22^ SFCs are constructed by recursive geometric procedure, which is schematically shown in Figure 1 for the Hilbert and Morton curves in the *d* = 2 case. The construction of the Hilbert curve proceeds by successively partitioning the square into subsquares and arranging them such that the curve passes through subsquares that share an edge. The same procedure is iteratively repeated for each subsquare. The Morton curve differs in the order of the traversal of the subsquares. The approximations to the two-dimensional SFC at finite resolution are the piecewise linear curves with 2^2*w*^ segments (approximating polygons), where *w* = 1, 2, … is the resolution index. The SFC results in the limit *w* → ∞. The construction can be generalized to *d* dimensions (*d* > 2), in which case the approximating polygon at resolution index *w* has 2^*dw*^ segments and traverses the entire (hyper)cube in ℝ^*d*^. By varying the relative orientations of the basic patterns and the approximating polygons in the recursion, a family of related *d*-dimensional SFCs of each curve type results for *d* > 2.^19^ The SFC encoding is reversible, that is, for every segment of the SFC, we can determine the corresponding location in the *d*-dimensional space.

**Figure 1.**
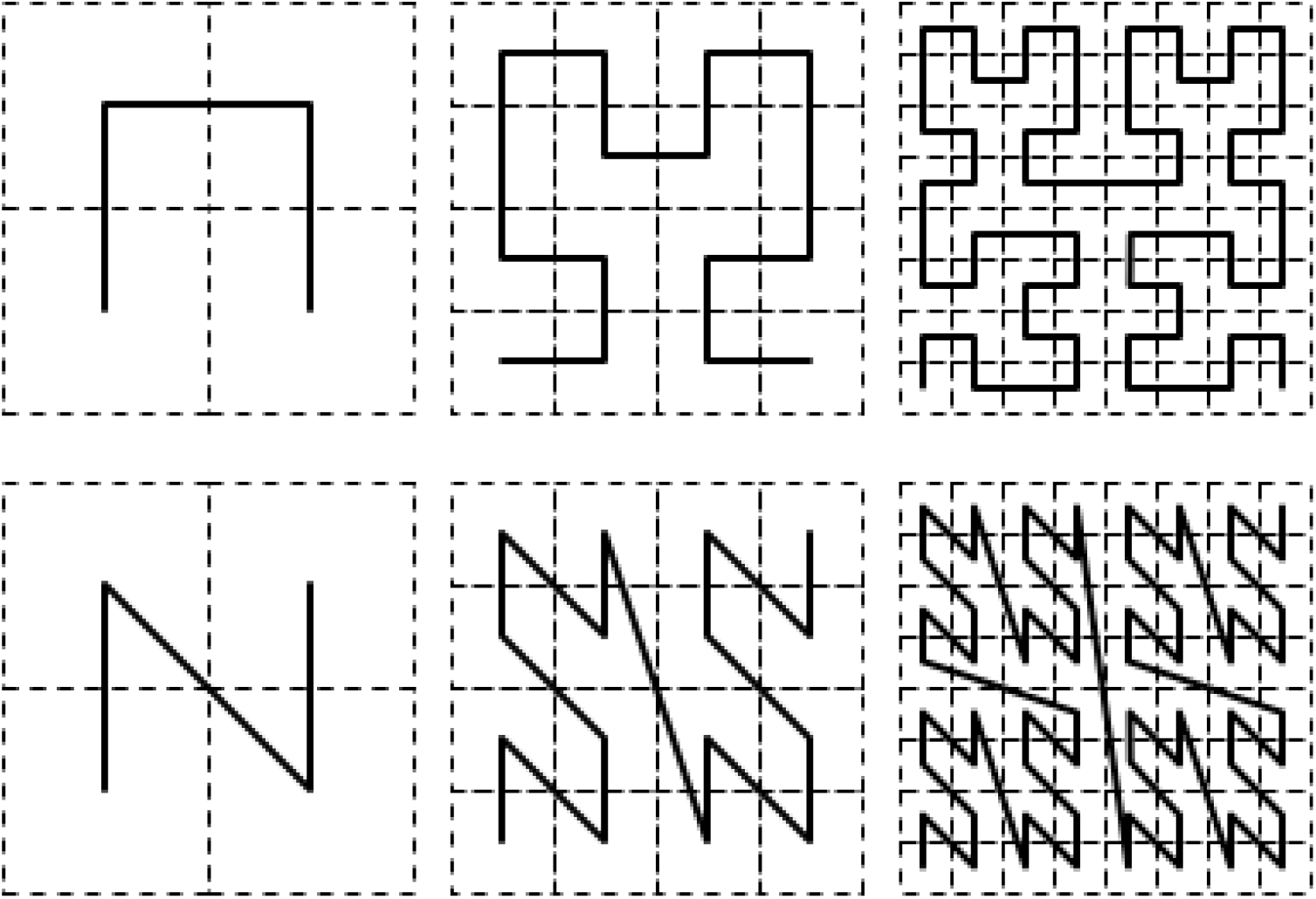
Recursive construction of two-dimensional Hilbert curve (top) and Morton Z-curve (bottom). Dashed lines indicate subsquare boundaries.

The self-similar construction of SFCs ensures that points that are close in the linear SFC ordering are mapped to near points in ℝ^*d*^. Conversely, most pairs of near points in ℝ^*d*^ are also close to each other in the SFC ordering. Due to this locality-preserving property, SFCs are a popular method for transforming multidimensional problems to one-dimensional ones with applications in optimization,^23–26^ multidimensional indexing,^27,28^ numerical simulations,^29,30^ and parallel algorithms.^31,32^ The locality preservation of the inverse mapping ℝ^*d*^ → [0, 1] is necessarily imperfect—one can always find a pair of points in ℝ^*d*^ that are far apart in the SFC ordering.^18,19^ Consider, for example, the turning points on the opposite sides of the two-dimensional Hilbert curve in Fig. 1. Locality measures corresponding to the average and worst-case distance in ℝ^*d*^ for neighboring points in the SFC ordering have been evaluated for different curve types and dimensionalities.^19,33,34^ The Hilbert curve is optimal with respect to the worst-case locality measure in the two-dimensional case and has been shown to perform well in practice. Considering these properties, SFCs are attractive as a minimally parameterized, readily generalizable encoding technique for three-dimensional structure data as input to structure prediction models.

As protein function can be described in multiple ways,^35^ we here focus on the enzyme-to-substrate mapping problem as a specific instance of the larger protein functional annotation challenge. Several structure-based ^36–38^ and/or sequence-based ^7,39–74^ approaches to enzyme function prediction have been reported. However, the large majority of these methods focus on predicting an enzyme’s Enzyme Commission (EC) number, a 4 digit number that maps enzymes to a hierarchical classification scheme. The information obtained from predictions at this level of classification is often already encoded in the protein domains found within a sequence,^75^ which are readily accessible in protein databases such as UniProt,^1,75^ even for orphan, poorly annotated proteins. For example, querying a recent enzymatic function prediction software ^39^ with the sequence for an orphan oxidoreductase in *Mycobacterium tuberculosis* (Rv3502c) as input, the output is the predicted EC class 1.1.1.-. This EC number corresponds (hierarchically) to: oxidoreductases (EC 1); acting on the CH-OH group of donors (EC 1.1); with nicotinamide adenine dinucleotide (NAD) or NADP as acceptor (EC 1.1.1). The two predictions reported by the first and third EC number digits (EC 1 and EC 1.1.1) are encoded in the fact that the protein belongs to the “Short-chain dehydrogenase/reductase SDR” family (IPR002347), as per its “Family and Domains” annotation in UniProt and InterPro. Furthermore, while the prediction encoded in the second EC number digit (EC 1.1: “acts on an alcohol or hydroxyl (CH-OH) donor group”) places constraints on the structure of the unknown native substrates, the number of possible chemical structures is still vast. Instead, what is often needed are one or several more granular predictions of a substrate’s chemical structure for enzymes within a protein family, as these can help generate testable hypotheses and guide downstream experimental efforts. Furthermore, despite the importance of functional predictions in enzymes, structured training and test data for machine learning models of enzymatic function are difficult to obtain.

We here first introduce a standardized dataset of annotated enzymes from two important protein families - the short-chain dehydrogenase/reductases (SDRs) and the S-adenosylmethionine-dependent methyltransferases (SAM-MTases) - and a set of labels that classify them according to their cofactor and substrate structural class preference. This dataset can be used to assess different protein representations and their use as input for machine learning methods that predict enzyme-substrate associations. We then show that a representation of protein structure using SFCs can be used to accurately predict enzyme cofactor and substrate structural class preference across a wide range of tasks. We find that orientational sampling of protein structural conformations increases the accuracy of both cofactor and substrate structural class predictions. The SFC-based representation of protein structure introduced here depends on significantly fewer parameters than commonly used alternatives, and our results demonstrate its potentially wide applicability for enzyme structure-function mapping tasks, and represent a baseline on which to build more refined models.

## Methods

### Structural Encoding

To generate feature representations of three-dimensional protein structures, the all-atom input coordinates were first preprocessed to remove translational and rotational degrees of freedom and coarse-grained at the amino acid residue level. Each residue was represented by its center of mass coordinates (computed using non-hydrogen atoms) and its type. The preprocessed coordinates were converted to the SFC-based feature representation by discretizing the space coordinates, SFC-based encoding, and subsequent binning. The resulting representation is a sparse fixed-length binary vector that is easy to use in machine learning tasks, for example, classification, regression, or similarity calculations. The structural encoding procedure is summarized in Table 2. All coordinates are in atomic units (au).

**Table 1.**
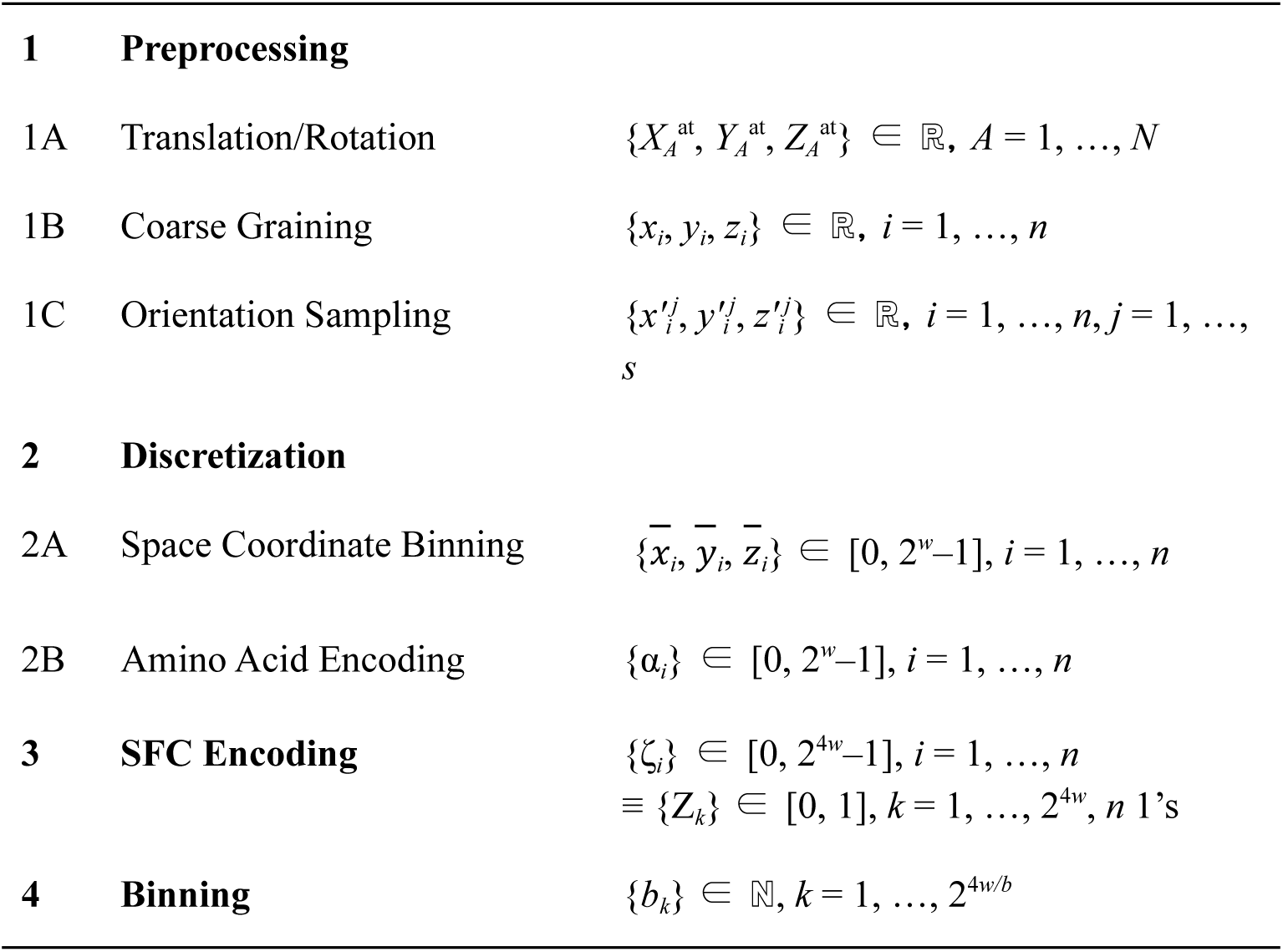
Scheme of protein three-dimensional structure encoding algorithm.

**Table 2.** Modified Li–Koehl AA encoding (5-bit).

| AA | $\alpha'$ | AA | $\alpha'$ | AA | $\alpha'$ | AA | $\alpha'$ |
| --- | --- | --- | --- | --- | --- | --- | --- |
| Ala | 21 | Gln | 25 | Leu | 0 | Ser | 22 |
| Arg | 26 | Glu | 24 | Lys | 17 | Thr | 23 |
| Asn | 31 | Gly | 29 | Met | 1 | Trp | 13 |
| Asp | 30 | His | 27 | Phe | 11 | Tyr | 10 |
| Cys | 7 | Ile | 2 | Pro | 20 | Val | 3 |

The translation/rotation preprocessing step (1A) removes the dependence of the input coordinates on arbitrary translations and rotations of the coordinate axes by transforming them to the structure’s inertial frame coordinate system. The coordinate origin is moved to the center of mass (computed using non-hydrogen atoms), and the coordinate axes are oriented along principal axes of inertia in the order of decreasing moment of inertia. The resulting all-atom coordinates 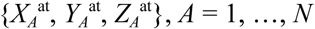, where *N* is the number of atoms, only depend on the internal degrees of freedom. We note that for a series of similar proteins, the use of inertial-frame coordinates improves the superposition of their three-dimensional structures but does not guarantee optimal alignment. The use of orientation sampling (see step 1C below) alleviates this issue, however, dedicated methods for multiple three-dimensional alignment should also be considered.^76^

The following coarse-graining step (1B) represents each amino acid residue by its center-of-mass coordinates (based on non-hydrogen atoms) {*x*_*i*_, *y*_*i*_, *z*_*i*_}, *i* = 1, …, *n*, where *n* is the number of amino acids, and its residue type. Coarse graining is convenient in protein studies^77^ because it encodes the internal structure of each residue as a single discrete variable (see 2B below) but all-atom approaches are more appropriate in other classes of molecules.

The exact SFC encoding can be approximated at different finite resolutions, which we identify by their bit width *w*. For a given value of *w*, each input coordinate (both spatial and residue type) is represented on a discrete grid of 2^*w*^ points. The continuous spatial coordinates are first discretized as integer indices 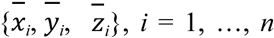, by placing the protein structure on a uniform cubic grid with 2^*w*^ cells per Cartesian direction and side length *L* (step 2A). The amino acid residue type is also encoded by an integer index {α_*i*_}, *i* = 1, …, *n*, in the range [0, 2^*w*^ – 1] using a procedure described in the next section (step 2B). The SFC encoding algorithm requires all coordinates to have the same number of grid points. The range of values for the residue type is thus rescaled if necessary to the same number of grid cells as the spatial coordinates.

The SFC-based encoding (step 3) converts each quadruple 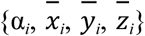 of *w*-bit integers losslessly to a 4*w*-bit integer coordinate *ζ*_*i*_ that gives its location index along the space-filling curve in the four-dimensional space. In this work, we use both the Hilbert SFC^18,21^ and the Morton Z curve^22^ for structure encoding. We note that the encoding is not invariant with the respect to the ordering of the Cartesian coordinates: the change in the first coordinate α_*i*_ is weighted more strongly than the changes in the subsequent coordinates 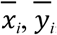, and 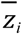 (in order of decreasing weight).

The SFC coordinate *ζ*_*i*_ indicates the presence of an amino acid residue of type α_*i*_ at the location given by the integer indices 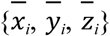. It follows that we can alternatively view the vector of *n* integer coordinates of 4*w* bit length as an extremely sparse binary vector {*Z*_*k*_}, *k* = 1, …, 2^4*w*^, which contains only *n* non-zero elements. This alternative view represents the protein structure as an unordered set of *n* locations (out of 2^4*w*^ possible) that are occupied by amino acid residues. This representation has several useful properties. It is that it is invariant with respect to a renumbering of the residues. Moreover, it enables comparisons between proteins of different length *n*. Finally, it can be used to construct feature representations at different resolutions by binning (see below). The mapping of the ordered vector {*ζ*_*i*_}, *i* = 1, …, *n*, to the binary vector (unordered set) {*Z*_*k*_} may seem to lose information about the order of amino acid residues in the protein sequence. However, this is not the case because the residue locations (and types) are encoded in the indices *ζ*_*i*_ and can be reconstructed when needed. Additionally, two different residues are extremely unlikely to share an index *ζ*_*i*_ unless the box size *L* is too small. In the absence of these improbable collisions, the vector {*ζ*_*i*_} and the binary vector {*Z*_*k*_} are equivalent.

Finally, the binning (step 4) compresses the vector *Z* by aggregating its entries in bins of width 2^*b*^. The resulting integer vector {*b*_*k*_} is of fixed length 2^4*w*/*b*^ with at most *n* non-zero elements. Due to the locality-preserving property of SFCs, each bin maps to a contiguous subset of the four-dimensional space, and consecutive bins correspond to neighboring subsets.

The structure encoding algorithm is implemented in the molz Python library and is released under the Apache 2 open-source license.^78^

### Amino Acid Encoding

Several numerical encodings of natural amino acids (AA) have been proposed, of which the most common ones are based on substitution frequencies in proteins ^79–85^ or the physical or conformational properties of amino acids.^86–94^ The first group of encodings is based on the evolutionary notion that amino acids are more likely to be homologous with similar amino acids. The amino acid encoding from the substitution frequency matrix^79,80,82,95^ by embedding in low-dimensional Euclidean spaces.^81,83,84^ The second group of encodings starts from an empirical set of physical and chemical properties of AAs and reduces them to low-dimensional vectors by principal component analysis (PCA).^87,92,94^

In this work, we used a discretized version of the five-dimensional encoding of Li and Koehl (LK), which was obtained from the BLOSUM62 similarity matrix by including the 5 largest principal components.^84^ To obtain an integer representation compatible with the discretized spatial coordinates, the LK feature vectors for the 20 natural AAs were arranged in the order of their decreasing PCA eigenvalues and encoded as integers α′ = (*a*_0_, *a*_1_, *a*_2_, *a*_3_, *a*_4_) in the range [0, 2^5^–1 = 31] with the *j*-th bit *a*_*j*_ = 0 if the *j*-th PCA component in the LK encoding was negative and 1 if it was positive. This approach generated several duplicates for similar amino acids, which were resolved manually, giving the encoding shown in Table 2. To generate a compatible grid to the spatial coordinates for SFC encoding, the 5-bit encodings α′ were uniformly scaled as α = 2^*w*–5^ α′.

Additionally, we investigated an encoding based on the five-dimensional z-scales of Wold and co-workers.^91,92^ The results are described in the supporting information (SI).

### Orientation Sampling

For a given pair of three-dimensional structures of the same composition, optimal alignment in terms of the root-mean-squared deviation can be efficiently computed.^96,97^ However, extensions to multiple alignment and different compositions are non-trivial. Instead of trying to find a single best alignment, we replicated the input protein structure under a fixed set of orientations 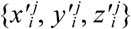, *i* = 1, …, *n, j* = 1, …, *s*, which are chosen to uniformly sample the space *SO*(3) of rotations in ℝ^3^. The underlying assumption in our approach is that, for any given pair of structures, one or more sampled orientations will be reasonably close to the optimal superposition. A deterministic and computationally simple approach to uniform sampling of rotations is the method of successive orthogonal images (SOI).^98^ The base grid generated by the SOI method consists of *s* = 72 orientations, which can be progressively refined. In this work, we limited ourselves to considering only the base SOI grid. The SFC encoding was carried out independently for each orientation to generate the binary vectors 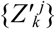, *j* = 1, …, *s*, which were subsequently combined into sparse binary vector 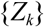, *k* = 1, …, 2^4*w*^ by binary OR operations. The resulting vector contains *ns* non-zero elements, barring collisions, and can be further aggregated to give the fixed-width integer vector {*b*_*k*_}, *k* = 1, …, 2^4*w*/*b*^, as in the base case.

### Enzyme selection and benchmark set

Enzymes belonging to the SDR and SAM-dependent methyltransferases were obtained from the UniProt database.^1^ Specifically, we performed queries using InterPro protein family/domain identifiers corresponding to each family: IPR002347 for short-chain dehydrogenase/reductase (SDRs) and IPR029063 for S-adenosylmethionine-dependent methyltransferases (SAM-MTases). The resulting tables contain each enzyme’s UniProt entry name, amino acid sequence and UniProt annotation score, a heuristic measure of annotation favoring literature-curated entries with experimental evidence.^99^ In addition for experimentally well-annotated enzymes, it includes Rhea^100^ and ChEBI^101^ database identifiers mapping each enzyme to the catalyzed biochemical reaction and its substrate and products structure represented as SMILES strings.^102^ We filtered the full set of enzymes in each family, keeping only those with UniProt annotation scores above “4-out-of-5”, to obtain the subset of well-annotated enzymes for inclusion in the benchmark set. The lists of protein structures were de-duplicated by their amino acid sequences, leaving 358 distinct SDR and 953 distinct SAM-MTase structures. The compiled benchmark data are included in the SI.

### Labeling enzymes using substrate clustering

Our first approach to classify enzymes according to the structural properties of the compounds they act on is based on structural clustering of the substrates and products of all annotated enzymes within a family. Broadly, we reasoned that one way to frame a machine learning classification task is to predict whether a given enzyme can catalytically act on substrates that belong to a structurally-related group of compounds, with the relevant clusters of compounds defined in an unsupervised manner. To implement this approach, we first removed all cofactors (NAD(H), NADP(H) for SDRs and S-adenosylmethionine for SAM-MTases), which appear in every reaction within each enzyme family, from the list of reaction components. Then, using RDKit,^103^ we took as input the SMILES string representations of all enzymatic substrates and products and obtained their corresponding Morgan fingerprint^104^ as bit vectors with radius = 3. We then generated a 2-dimensional projection of the Morgan fingerprints using the UMAP algorithm,^105^ with the Jaccard metric for binary vectors. Since we eventually trained separate machine learning classifiers for each enzyme family, we note that the SDR and SAM-MTase substrates and products were processed separately at this stage. Finally, using *k*-means clustering ^106,107^ we clustered the compounds according to their 2D UMAP projection,^105^ choosing the optimal number of clusters that maximized the silhouette score.^108^ This clustering procedure generated 9 structure clusters for SDR substrates (numbered 0–8, as shown in Figure 2) and 13 structure clusters for SAM-MTase substrates (numbered 0–12). The substrate structures and enzyme clusters are included in the SI.

**Figure 2.**
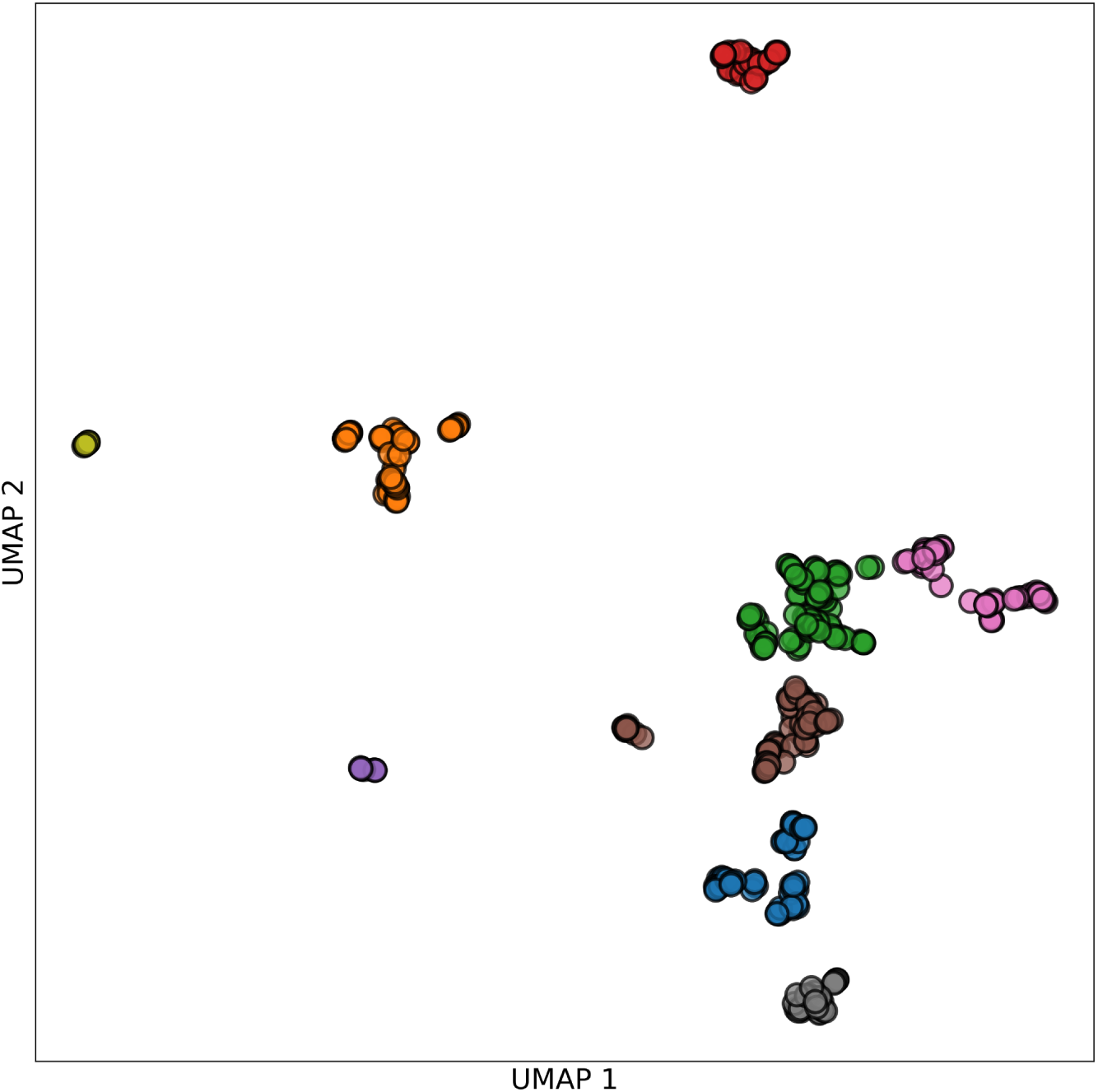
Structural clustering of substrates and products of annotated short-chain dehydrogenase / reductase (SDR) enzymes. Each circle represents a unique substrate or product. These compounds were clustered according to their chemical structure by (1) obtaining Morgan fingerprints from their SMILES representation; (2) projecting the Morgan fingerprints to 2-dimensions using UMAP; and (3) grouping compounds using k-means clustering.

### Labeling enzymes using substructure search on substrates and products

Our second approach to label enzymes according to the type of compound they act on is based on searching for broad classes of substrates represented by molecular substructures. We used RDKit to search for molecular patterns (phenol, sterol, or acyl-CoA) encoded as SMARTS strings.^109^ The presence or absence of these substructures in the substrates or products was treated directly as a classification label. The number of SAM-MTases acting on sterols was too small for making statistically significant predictions. Thus we did not consider this classification task further.

## Results

### Benchmark Datasets

Our standardized enzymes-substrate dataset contains reference data for 358 unique short-chain dehydrogenase reductases (SDRs) and 953 unique S-adenosylmethionine dependent methyltransferases (SAM-MTases) structures. For SDRs, the reference data includes the preference for the redox cofactor (NAD(H) or NADP(H)) and the specificity with respect to a broad substrate class (e.g phenols, sterols, and acyl-CoA substrates). For SAM-MTases, the enzymes acting on biopolymers (proteins or nucleic acids) or on small molecules are distinguished. The enzyme specificity with respect to two substrate classes (phenols and N-heterocycles) is also classified. Additionally, the binary classifications of substrate specificities of SDRs and SAM-MTases by substrate clusters from unsupervised clustering are included in the benchmark set.

We constructed three-dimensional feature representations of 358 unique SDR and 953 unique SAM-MTase structures and assessed their performance on a set of binary classification tasks related to their cofactor and substrate specificity. The initial all-atom structures were generated by the public version of the AlphaFold2 protein folding approach^4^ via the ColabFold notebook.^110^ MMSeq2 multiple sequence alignment algorithm ^111,112^ was used instead of the original AlphaFold2 homology search. The protein sequences in the FASTA format and the three-dimensional protein structures in the PDB format are included in the SI.

The performance of the three-dimensional feature representations was assessed in enzymatic function predictions using our benchmark set. For SDRs, the benchmark set contains binary classification tasks with respect to the redox cofactor (NAD(H) or NADP(H)) and to substrate class (phenols, sterols, and acyl-CoA). For SAM-MTases, the classification of enzymes acting on biopolymers (proteins or nucleic acids) or on small molecules and substrate specificity with respect to phenols and N-heterocycles are included in the benchmark set. Additionally, binary classifications of enzyme specificity of SDRs and SMT-MTases based on substrate clusters from unsupervised clustering are part of the benchmark set. The protein structures were preprocessed and converted to three-dimensional feature representations as described in the Methods section. Both Hilbert and Morton SFCs were studied for the SFC encoding. The discretization box was of side length *L* = 200 au (100 au for SAM-MTases) and of *w* = 8 bit width resolution. The binary vectors were binned using 2^*b*^ = 256, 4096, and 65536 bins. Orientation sampling using the SOI method at the base grid level (*s* = 72 orientations) was applied. In the following, we evaluate the performance of three-dimensional feature representations as a function of the classification task, SFC type, and orientation sampling. We compare the full encodings containing the three-dimensional protein structure representations including the residue types with simplified encodings containing only amino acid types (bag-of-amino acids representation) or only the residue positions, regardless of amino acid type (structure-only representation).

The classifications were performed using the XGBoost binary classifier with GBTree booster, maximum depth 8, learning rate 0.2, and *L*_2_ = 1 regularization.^113^ For each classification task, we report the 5-fold cross validated accuracy (percentage of correct predictions) and the receiver operator characteristic (ROC) area under curve (AUC). All classifications were performed using the scikit-learn library, version 1.0.2.114 The results of hyperparameter optimization using Bayesian optimization methods are included in the SI. We find that the results are not strongly dependent on the parameter choice. Classifications using support vector machine classification (SVMC)^115^ and LightGBM methods^116^ produced similar results and are reported in the SI.

### Classification Task

The accuracy and AUC characteristics of 4 binary classification tasks for SDRs and 3 binary classification tasks for SAM-MTases using Hilbert space-filling curve are given in Table 3. Figure 3 shows the ROC curve for the NAD/NADP classification with and without random shuffling of the labels. We obtain accuracy values (percentage of correctly predicted classes) between 0.766–0.906 for the classification tasks. The AUC characteristics for these binary classification tasks are between 0.828–0.922. The classification performed using randomly shuffled labels shown in Figure 1 serves as a null hypothesis and predicts the AUC close to the theoretically expected value of 0.5 for a random binary classifier.

**Table 3.** Accuracy and ROC area under curve (AUC) of structure encodings of SDRs and SAM-MTases using 8-bit Hilbert SFC, modified LK encoding (Table 2), 4096 bins, orientation sampling using SOI (s = 72) for binary classification tasks (5-fold cross validated).

| SDR | Accuracy | AUC | SAM | Accuracy | AUC |
| --- | --- | --- | --- | --- | --- |
| NAD/NADP | 0.766 | 0.837 | Biopolymer/<br>small molecule | 0.848 | 0.922 |
| Phenols | 0.809 | 0.828 | Phenols | 0.829 | 0.876 |
| Sterols | 0.832 | 0.833 | N-Heterocycles | 0.787 | 0.842 |
| Acyl-CoA | 0.906 | 0.900 |  |  |  |

**Figure 3.**
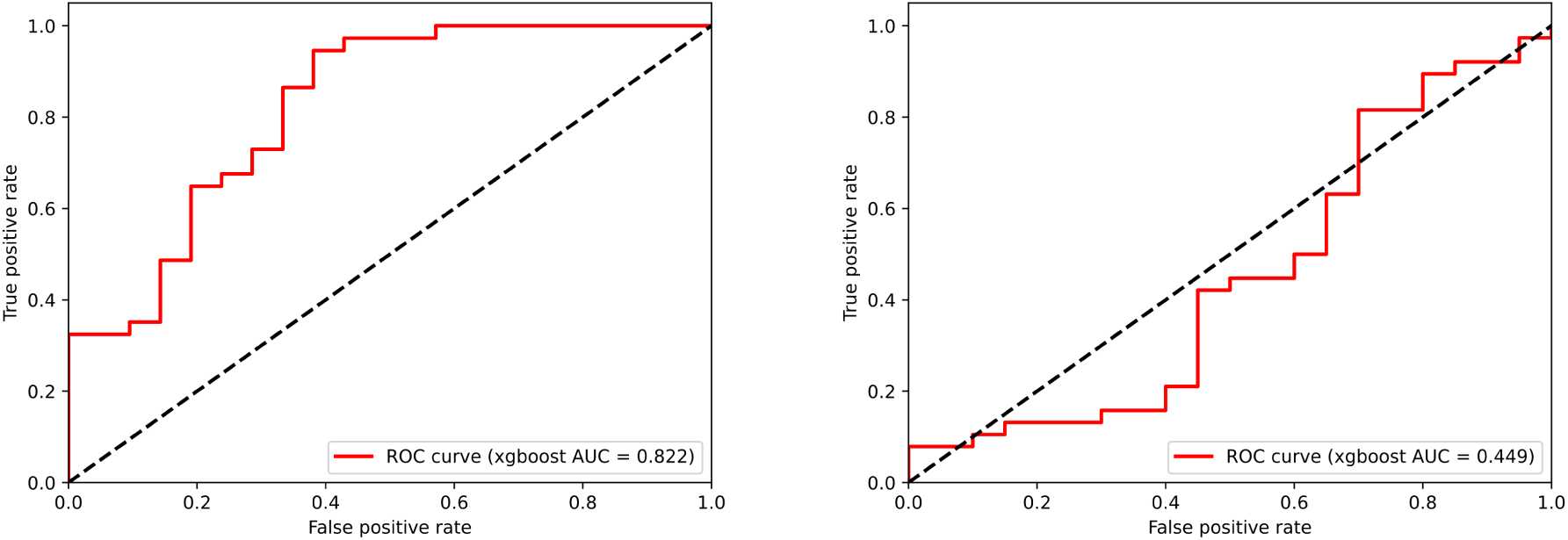
ROC curves for NAD/NADP classification of SDRs using 8-bit Hilbert SFC with modified LK encoding, 4096 bins, orientation sampling using SOI (s = 72, left), same after random label shuffle (right).

The results of binary classification tasks for substrate specificity to substrate clusters obtained from unsupervised clustering are shown in Table 4. Only clusters including 5 or more molecules (6 clusters for SDRs and 7 clusters for SAM-MTases) were included in classification tasks. The accuracy values of classification tasks with respect to clusters are in the range of 0.728–0.942 and the AUC values are 0.717–0.975. We note that the classification results are similar for SDRs and SAM-MTases.

**Table 4.** Accuracy and ROC area under curve (AUC) of structure encodings of SDRs and SAM-MTases using 8-bit Hilbert SFC, modified LK encoding (Table 2), 4096 bins, orientation sampling using SOI (s = 72) for binary classification tasks from unsupervised clustering (5-fold cross validated). Only tasks with at least 5 examples per class were considered.

| SDR | Accuracy | AUC | SAM | Accuracy | AUC |
| --- | --- | --- | --- | --- | --- |
| Cluster 0 | 0.926 | 0.754 | Cluster 0 | 0.864 | 0.878 |
| Cluster 1 | 0.896 | 0.852 | Cluster 1 | 0.942 | 0.908 |
| Cluster 2 | 0.815 | 0.837 | Cluster 5 | 0.876 | 0.717 |
| Cluster 4 | 0.864 | 0.835 | Cluster 6 | 0.915 | 0.889 |
| Cluster 5 | 0.945 | 0.975 | Cluster 7 | 0.892 | 0.837 |
| Cluster 7 | 0.728 | 0.763 | Cluster 8 | 0.934 | 0.963 |
|  |  |  | Cluster 11 | 0.880 | 0.917 |

### Space-Filling Curve Type

Using Morton Z curve instead of Hilbert SFC for the encoding gave very similar binary classification results, which are shown in Table 5. The accuracy values were in the range of 0.762–0.851, while the AUC values were between 0.825–0.920, almost all of them within one percent point of the results obtained with Hilbert SFC.

**Table 5.** Accuracy and ROC area under curve (AUC) of structure encodings of SDRs and SAM-MTases using 8-bit Morton SFC, modified LK encoding (Table 2), 4096 bins, orientation sampling using SOI (s = 72) for binary classification tasks (5-fold cross validated).

| SDR | Accuracy | AUC | SAM | Accuracy | AUC |
| --- | --- | --- | --- | --- | --- |
| NAD/NADP | 0.762 | 0.844 | Biopolymer/<br>small molecule | 0.851 | 0.920 |
| Phenols | 0.825 | 0.825 | Phenols | 0.833 | 0.880 |
| Sterols | 0.828 | 0.828 | N-Heterocycles | 0.794 | 0.845 |
| Acyl-CoA | 0.899 | 0.900 |  |  |  |

### Bag-of-Amino Acids and Structure-Only Representations

As described above, the SFC encoding imposes an ordering in the *d*-dimensional representation in that a change in the first coordinate of the *d*-dimensional representation induces a larger difference in the SFC ordering than a change in the second coordinate, etc. Since we apply the SFC encoding to the tuple 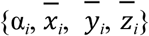, the residue type α_*i*_ has by design the largest weight in the encoded vector. In fact, in the limit of an infinitely large discretization box (side length *L*→∞), all atomic positions are mapped to the origin and only the residue type information remains. The SFC encoded vector approaches a very simple model of the protein structure, which can be denoted as a bag-of-amino acids. This representation contains only the counts of residue types and no spatial or sequence information. Interestingly, this feature representation produces quite similar results to the full SFC encoding, as shown in Table 7. The accuracy values for the binary classification tasks were between 0.806–0.909 and the AUC values were found to be in the range 0.788–0.924.

**Table 7.** Accuracy and ROC area under curve (AUC) of bag-of-amino acids encodings of SDRs and SAM-MTases using modified LK encoding (Table 2), 32 bins for binary classification tasks (5-fold cross validated).

| SDR | Accuracy | AUC | SAM | Accuracy | AUC |
| --- | --- | --- | --- | --- | --- |
| NAD/NADP | 0.808 | 0.884 | Biopolymer/<br>small molecule | 0.844 | 0.924 |
| Phenols | 0.819 | 0.856 | Phenols | 0.806 | 0.878 |
| Sterols | 0.861 | 0.863 | N-Heterocycles | 0.748 | 0.788 |
| Acyl-CoA | 0.909 | 0.864 |  |  |  |

To test the importance of the residue type and spatial information, we also considered the converse case of the structure-only encoding, which contains only the three-dimensional coordinates of the residue but no residue type information. The results are shown in Table 8 and again perform similarly to those obtained with the full SFC encoding. The accuracy values were between 0.767–0.864 and the AUC values were 0.733–0.916.

**Table 8.**
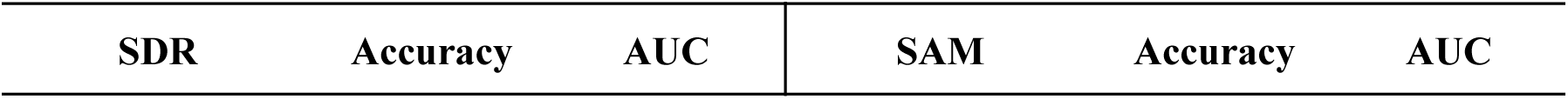
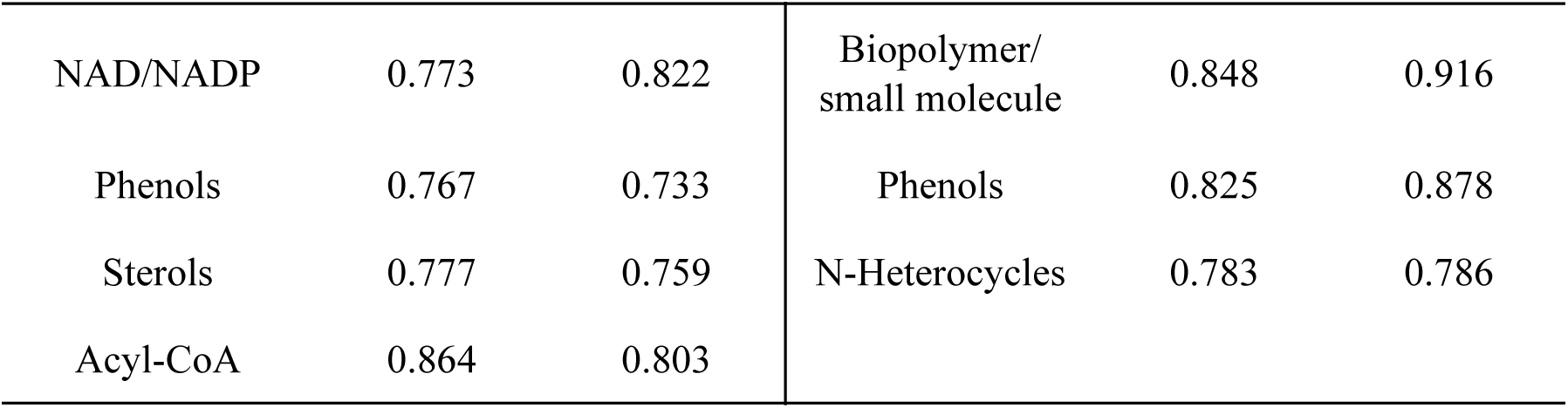
Accuracy and ROC area under curve (AUC) of structure-only encodings of SDRs and SAM-MTases using 8-bit Hilbert SFC, 4096 bins, orientation sampling using SOI (s = 72) for binary classification tasks (5-fold cross validated).

## Discussion

As our results show, enzymatic function can be predicted from feature representations based on the three-dimensional structures of SDRs and SAM-MTAses with good accuracy (> 80% binary classification accuracy, > 0.8 AUC). Moreover, the predictive power is not strongly dependent on the parameters of the representation, for example, type of the SFC encoding. As we show in the SI, other parameters such as the amino acid encoding and the number of bins do not affect the accuracy of the predictions. The weak variation of the results with the parameters and between two unrelated protein families is consistent with our construction, which includes relatively few adjustable “knobs” by design. As a result, the methodology described here should generalize well to other protein families and other functional prediction tasks. Our results are also encouraging for the modeling strategy which relies on minimally parameterized feature representations of protein structure. The predictions in this work were made with the XGBoost classification model, which has been shown to generalize well and to be relatively insensitive to model parameters. In the SI, we show the results of hyperparameter optimization of the XGBoost model for the NAD/NADP cofactor specificity of SDRs. Interestingly, the results obtained using optimized hyperparameters do not differ strongly from the parameter set used through this work. The binary classifications using the SVMC and LightGBM models also give similar accuracy, as shown in the SI. These findings support our assumption that the predictions are not strongly parameter-dependent.

The results obtained without orientational sampling are slightly worse, which suggests that structure alignment is important to the prediction. However, the differences between the results obtained with and without orientational sampling are not very large. However, global structural alignment may not be optimal for proteins with very different compositions, while methods for aligning active site regions may produce further improvements.

The fidelity of the 3D to 1D mapping can be analyzed in terms of the correlation between the inter-residue distances in three-dimensional space and along the SFC ordering. The average Pearson correlation coefficients for the Euclidean inter-residue distances and their encoding using the Hilbert SFC are *R*^2^ = 0.446 for SDRs and 0.465 for SAM-MTases. For the encoding using the Morton Z curve, the correlation coefficients are similar, giving *R*^2^ = 0.434 for SDRs and 0.495 for SAM-MTases. The correlation is affected by its dependence on the ordering of the *x, y, z* Cartesian components, in that the influence of the change in the Cartesian coordinates decreases with *x* > *y* > *z*. Orientation-invariant encodings might be helpful in addressing this issue.

The greatest surprise in our results was a significant amount of redundancy contained in the feature representations. The bag-of-amino acids representation, which consists of the counts of different amino acid types, yields predictions that are quite similar to the full feature representation containing both amino acid types and their spatial locations. On the other hand, the structure-only representation, which elides the amino acid types and only shows the presence of some amino acid at a location in space, also produces predictions with comparable accuracy. Two possible explanations may be responsible for this behavior. First, the sequence and the corresponding three-dimensional structure may be more strongly correlated with respect to the protein function than thus far appreciated. The correlations seem to be present within each SDR and SAM-MTase protein families. This can be understood since the three-dimensional structures are obtained by applying the AlphaFold2 model to the protein sequence. While this work is focused on naturally occurring protein structures, it might be useful to consider synthetic data sets with a lesser degree of correlation between the structure and sequence in order to explore this phenomenon. Second, the uniform discretization of the three-dimensional structure might be insufficiently sensitive to the active site structure, which would require finer resolution. In order to better explore this hypothesis, multiscale features representation should be explored, which are not based on an encoding of a regular grid in 3D (or 4D) space but instead on an adaptive-resolution grid.

The design of methods for protein function predictions is constrained by two somewhat clashing requirements. On one hand, only a fraction of the total protein structure constitutes its active site and is involved in its function. The remainder of the sequence may provide the crucial structural scaffold but its structure may provide little discriminating power with respect to the enzyme function. Relatively large structural changes outside of the active site may be functionally of little consequence. On the other hand, relatively small changes in the structure of the active site can be expected to have a significant effect on the enzyme function. Therefore, the function prediction methods have to simplify the structural parameters away from the active site, e.g., by graph models or discretization, while remaining sensitive to the structure of the active site. This supports the need for encodings with adaptive resolution.

The advantage of SFC-based encodings is that they are discrete, reversible, and deterministic. As a result, generative models can be formulated in the SFC encoding, which are straightforward to transform back to the full 3D (or 4D) coordinates. On the other hand, this means that the preservation of locality is only approximate and the worst-case behavior is effectively unbounded, as discussed in the introduction. If one is willing to give up the deterministic construction, random projection models can give better average or worst-case behavior.

## Supporting Information

### Amino Acid Encoding

We investigated an additional AA encoding based on the five-dimensional z-scales of Wold and co-workers.^91,92^ Unlike the LK encoding, which is derived from substitution matrices, the z-scales are based on physical and electronic properties of amino acids. The five-dimensional z-scale encoding^92^ was obtained from dimension reduction of a set of 26 AA descriptors using PCA. After applying the discretization approach as for the LK encoding and manually resolving duplicates, we arrived at the AA encoding shown in Table S1. Surprisingly, the two AA encodings are quite similar to each other, despite differences in derivation.

**Table S1.** Modified z-scale AA encoding (5-bit).

| AA | $\alpha'$ | AA | $\alpha'$ | AA | $\alpha'$ | AA | $\alpha'$ |
| --- | --- | --- | --- | --- | --- | --- | --- |
| Ala | 20 | Gln | 25 | Leu | 2 | Ser | 21 |
| Arg | 26 | Glu | 24 | Lys | 27 | Thr | 16 |
| Asn | 29 | Gly | 19 | Met | 6 | Trp | 12 |
| Asp | 28 | His | 31 | Phe | 14 | Tyr | 13 |
| Cys | 22 | Ile | 1 | Pro | 15 | Val | 0 |

**Table S2.** Accuracy and ROC area under curve (AUC) of structure encodings of SDRs and SAM-MTases using 8-bit Hilbert SFC, modified z scale encoding (Table 3), 4096 bins, orientation sampling using SOI (s = 72) for binary classification tasks (5-fold cross validated).

| SDR | Accuracy | AUC |
| --- | --- | --- |
| NAD/NADP | 0.755 | 0.812 |
| Phenols | 0.793 | 0.660 |
| Sterols | 0.770 | 0.760 |
| Acyl-CoA | 0.893 | 0.823 |

**Table S3.**
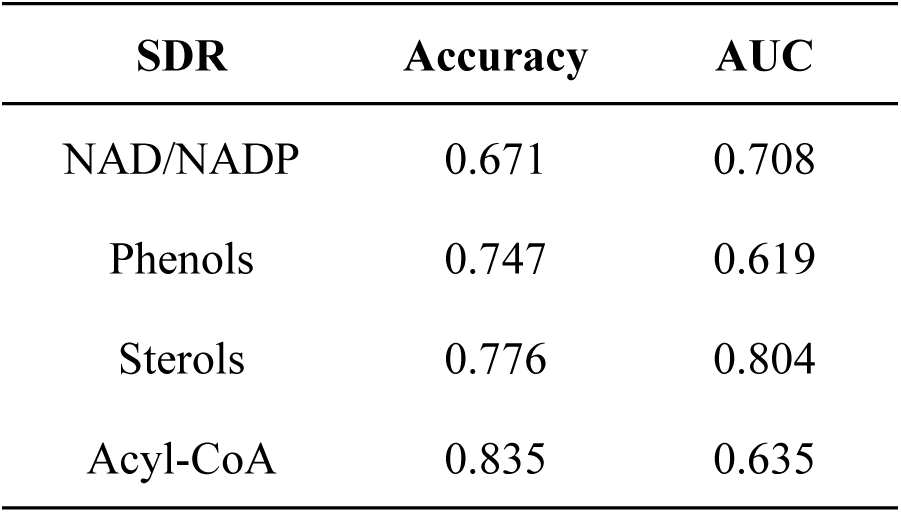
Accuracy and ROC area under curve (AUC) of bag-of-amino acids encodings of SDRs and SAM-MTases using modified z scale encoding (Table 3), 32 bins for binary classification tasks (5-fold cross validated).

### Orientation Sampling Using SOI

Results without orientation sampling are shown in Table S4.

**Table S4.** Accuracy and ROC area under curve (AUC) of structure encodings of SDRs and SAM-MTases using 8-bit Hilbert SFC, modified LK encoding (Table 2), 4096 bins for binary classification tasks (5-fold cross validated).

| SDR | Accuracy | AUC | SAM | Accuracy | AUC |
| --- | --- | --- | --- | --- | --- |
| NAD/NADP | 0.738 | 0.782 | Biopolymer/<br>small molecule | 0.822 | 0.907 |
| Phenols | 0.799 | 0.637 | Phenols | 0.794 | 0.841 |
| Sterols | 0.786 | 0.700 | N-Heterocycles | 0.682 | 0.728 |
| Acyl-CoA | 0.887 | 0.757 |  |  |  |

### Binning

**Table S5.** Accuracy and ROC area under curve (AUC) of structure encodings of SDRs and SAM-MTases using 8-bit Hilbert SFC, modified LK encoding (Table 2), 256 bins, orientation sampling using SOI (s = 72) for binary classification tasks (5-fold cross validated).

| SDR | Accuracy | AUC | SAM | Accuracy | AUC |
| --- | --- | --- | --- | --- | --- |
| NAD/NADP | 0.699 | 0.756 | Biopolymer/<br>small molecule | 0.816 | 0.894 |
| Phenols | 0.816 | 0.805 | Phenols | 0.813 | 0.881 |
| Sterols | 0.816 | 0.831 | N-Heterocycles | 0.744 | 0.778 |
| Acyl-CoA | 0.899 | 0.799 |  |  |  |

**Table S6.** Accuracy and ROC area under curve (AUC) of structure encodings of SDRs and SAM-MTases using 8-bit Hilbert SFC, modified LK encoding (Table 2), 65536 bins, orientation sampling using SOI (s = 72) for binary classification tasks (5-fold cross validated).

| SDR | Accuracy | AUC | SAM | Accuracy | AUC |
| --- | --- | --- | --- | --- | --- |
| NAD/NADP | 0.815 | 0.897 | Biopolymer/<br>small molecule | 0.865 | 0.942 |
| Phenols | 0.854 | 0.897 | Phenols | 0.825 | 0.878 |
| Sterols | 0.867 | 0.894 | N-Heterocycles | 0.771 | 0.824 |
| Acyl-CoA | 0.900 | 0.907 |  |  |  |

### Classification Algorithm

**Table S7.** Accuracy and ROC area under curve (AUC) of structure encodings of SDRs and SAM-MTases with SVM classification using 8-bit Hilbert SFC, modified LK encoding (Table 2), 4096 bins, orientation sampling using SOI (s = 72) for binary classification tasks (5-fold cross validated).

| SDR | Accuracy | AUC | SAM | Accuracy | AUC |
| --- | --- | --- | --- | --- | --- |
| NAD/NADP | 0.713 | 0.782 | Biopolymer/<br>small molecule | 0.846 | 0.912 |
| Phenols | 0.773 | 0.695 | Phenols | 0.810 | 0.873 |
| Sterols | 0.796 | 0.716 | N-Heterocycles | 0.798 | 0.828 |
| Acyl-CoA | 0.896 | 0.836 |  |  |  |

**Table S8.** Accuracy and ROC area under curve (AUC) of structure encodings of SDRs and SAM-MTases with LightGBM classification using 8-bit Hilbert SFC, modified LK encoding (Table 2), 4096 bins, orientation sampling using SOI (s = 72) for binary classification tasks (5-fold cross validated).

| SDR | Accuracy | AUC | SAM | Accuracy | AUC |
| --- | --- | --- | --- | --- | --- |
| NAD/NADP | 0.766 | 0.828 | Biopolymer/<br>small molecule | 0.857 | 0.932 |
| Phenols | 0.819 | 0.847 | Phenols | 0.837 | 0.871 |
| Sterols | 0.832 | 0.839 | N-Heterocycles | 0.787 | 0.843 |
| Acyl-CoA | 0.900 | 0.918 |  |  |  |

### Training and Test Set Size

**Table S9.** Training and test set size for classification tasks used in this work (5-fold cross validated).

| SDR | Training | Test | SAM | Training | Test |
| --- | --- | --- | --- | --- | --- |
| NAD/NADP | 228 | 58 | Biopolymer/<br>small molecule | 762 | 191 |
| Phenols | 247 | 62 | Phenols | 206 | 52 |
| Sterols | 247 | 62 | N-Heterocycles | 206 | 52 |
| Acyl-CoA | 247 | 62 | Clustering | 206 | 52 |
| Clustering | 247 | 62 |  |  |  |

### Hyperparameter Optimization

**Table S10.** Results of Bayesian hyperparameter optimization using accuracy and ROC area under curve (AUC) for the NAD/NADP classification task in SDRs using 8-bit Hilbert SFC, modified LK encoding (Table 2), 4096 bins

| **Objective** | **Best value** | **Learning Rate** | **Max Depth** | **$L_2$ reg.** | **$L_1$ reg.** |
| --- | --- | --- | --- | --- | --- |
| Accuracy | 0.825 | 0.031 | 16 | 2.172 | 0.271 |
| AUC | 0.868 | 0.078 | 14 | 1.318 | 0.058 |

SDR NAD/NADP classification, hyperparameter optimization for XGBoost algorithm using Bayesian optimization: learning rate (lognormal prior with log mu = 0, sigma = 1), max depth (flat discrete prior: 8, 10, 12, 14, 16), regularization alpha (L2, lognormal prior with log mu = 0, sigma = 1), regularization lambda (L1, lognormal prior with log mu = 0, sigma = 1). The results are given in Table S10. Note that the best accuracy and AUC values obtained are close to the values in Table 3.

## TOC Graphic

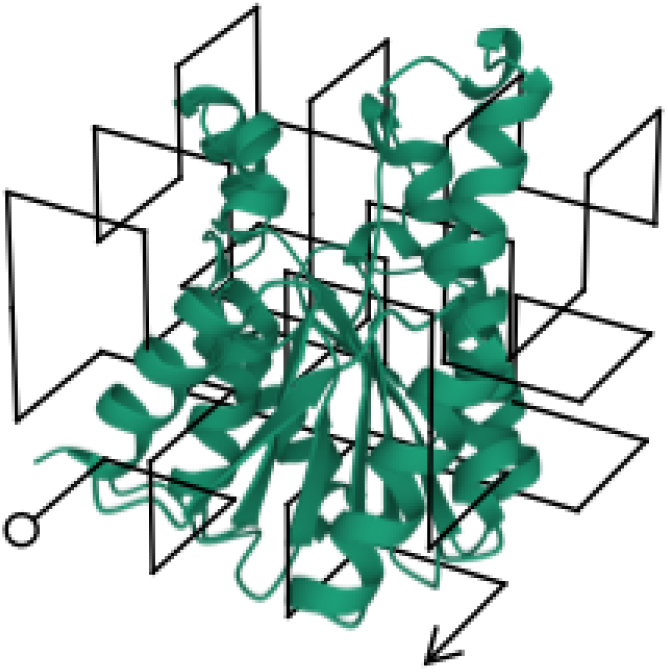

Three-dimensional model of the Short-chain dehydrogenase/reductase SDRA of *Arabidopsis thaliana* (Uniprot code: Q9S9W2) overlaid with the model of the three-dimensional Hilbert curve.

